# AOP-helpFinder 3.0: from text mining to network visualization of key event relationships, and knowledge integration from multi-sources

**DOI:** 10.1101/2025.04.22.648318

**Authors:** Thomas Jaylet, Florence Jornod, Quentin Capdet, Olivier Armant, Karine Audouze

## Abstract

**Motivation:** The Adverse Outcome Pathways (AOP) framework advances alternative toxicology by prioritizing the mechanisms underlying toxic effects. It organizes existing knowledge in a structured way, tracing the progression from the initial perturbation of a molecular event—caused by various stressors—through key events (KEs) across different biological levels, ultimately leading to adverse outcomes that affect human health and ecosystems. However, the increasing volume of toxicological data presents a significant challenge for integrating all available knowledge effectively.

**Results:** Artificial intelligence provides powerful methods to analyze and integrate large, heterogeneous data sources. Within this framework, the AOP-helpFinder text mining tool, accessible as a web server, was designed to identify stressor-event and event-event relationships by automatically screening scientific literature in the PubMed database, facilitating the development of AOPs. The proposed new version introduces enhanced functionality by incorporating additional data sources, automatically annotating events from the literature with toxicological database information in a systems biology context. Users can now visualize results as interactive networks directly on the web server. With these advancements, AOP-helpFinder 3.0 offers a robust solution for integrative and predictive toxicology, as demonstrated in a case study exploring toxicological mechanisms associated with radon exposure.

**Availability:** AOP-helpFinder is available at https://aop-helpfinder-v3.u-paris-sciences.fr/

**Supplementary information:** Supplementary data are available on Zenodo (https://zenodo.org/records/15193936) and codes on GitHub (https://github.com/systox1124/AOP-helpFinder).

## 1 Introduction

Human beings and ecosystems are exposed daily to an increasing number of potentially toxic substances and other factors (stressors), whose effects and risks are often poorly characterized, partly due to the low efficiency of traditional in vivo testing. Modern toxicology relies on new alternative approaches (NAMs) and high-throughput methodologies to address this limitation (Krewski et al. 2010, Krewski et al. 2020). In 2010, the concept of Adverse Outcome Pathways (AOPs), proposed by Ankley et al., emerged as an effective solution to integrate and structure existing toxicological knowledge, thus supporting mechanistic understanding and risk assessment in response to the growing demand for the evaluation of stressors (Ankley et al. 2010). An AOP describes a sequence of events that starts with a Molecular Initiating Event (MIE), triggered by a prototypical stressor (which is not part of the AOP itself). This MIE is linked to a sequence of key events (KEs) across different levels of biological organization, ultimately leading to an outcome (AO) at the individual or population level. These events (MIEs, KEs, AOs) are connected by key event relationships (KERs) that describe their causal links, enhancing the understanding of stressor(s) toxicity. Biological events can be shared across multiple AOPs, allowing the formation of Adverse Outcome Pathways Networks (AOPNs) that reflect the complexity of biological systems (Villeneuve et al. 2014, Yarar and Wojewodzic 2024).

Given the vast amount of available data, identifying and integrating relevant toxicological information into an AOP can be difficult and time-consuming. Artificial intelligence (AI) offers techniques to address these issues, and the AOP-helpFinder tool was developed with this objective in mind. Available as a web server, AOP-helpFinder uses text mining (TM), graph theory, and Natural Language Processing (NLP) methods to identify relationships between stressors and biological events (MIE, KE, AO), as well as between pairs of biological events (KE-KE), by analyzing PubMed abstracts and providing confidence scores for each link, therefore supporting weight of evidence (WoE) (Carvaillo et al. 2019, Jornod et al. 2022, Jaylet et al. 2023).

However, it is crucial to consider data from multiple sources, such as toxicological databases to cross AOPs with more existing knowledge. Therefore, we present an enhanced version of AOP-helpFinder that automatically links and annotates extracted information from the literature to various relevant databases using a systems biology approach. This version also includes the capacity to visualize the identified KER as interactive biological networks directly from the web server. A case study is presented to demonstrate the tool’s effectiveness, focusing on identifying and capturing potential mechanisms and pathologies induced after radon exposure, integrating scientific literature and multi-source databases knowledge.

## 2 Materials & Methods

### 2.1 Previous versions

The initial version of the AOP-helpFinder method, developed in 2019, was designed to accelerate the identification of associations between environmental stressors and biological events implicated in AOPs (Carvaillo et al. 2019). Given the rapid expansion of scientific literature within the PubMed database, it has become increasingly feasible to investigate potential links between stressors and biological events, whether catalogued in repositories such as AOP-Wiki or identified by domain experts. In response to the strong interest expressed by both scientific and regulatory communities, a web-based interface for AOP-helpFinder was developed and published (Jornod et al. 2022). The tool has since been applied in multiple studies, more than ten peer-reviewed publications to date, with additional research ongoing, and has contributed to the development of several AOPs (e.g., IDs 439, 441 & 490) submitted to the AOP-Wiki. Its utility has been recognized internationally in the field of toxicology, and it has recently been incorporated as a third-party tool within the OECD AOP-Wiki platform. Recently, an updated version of the method and the online platform (AOP-helpFinder 2.0) was created, that introduces several enhancements, the most notable being the implementation of an automated screening system for PubMed abstracts to detect and extract associations between biological events, including MIE, KE and AO (Jaylet et al. 2023). Furthermore, a refined scoring system has been developed to classify co-occurring terms— either stressor-event or event-event—into five priority categories. This classification is intended to support data prioritization and reinforce the weight-of-evidence framework essential for AOP development. To further facilitate the interpretation of results, new visualization features have also been integrated into the platform.

Recognizing that comprehensive AOP development requires integration of data beyond the literature, the proposed third version includes automated annotation of key biological events (MIE, KE, AO) with supplementary information from open-access toxicological databases (descriptions are on section 2.2 to 2.4, and all details can be found on the supplementary material https://zenodo.org/records/15193936). The AOP-helpFinder 3.0 tool is freely available, with all its informatics source code are accessible via GitHub (https://github.com/systox1124/AOP-helpFinder). A user-friendly web interface enables direct execution of literature searches & event annotations to support AOP development (https://aop-helpfinder-v3.u-paris-sciences.fr/). A complete workflow of the procedure is shown in Figure 1 to facilitate user understanding and usage.

**Figure 1.**
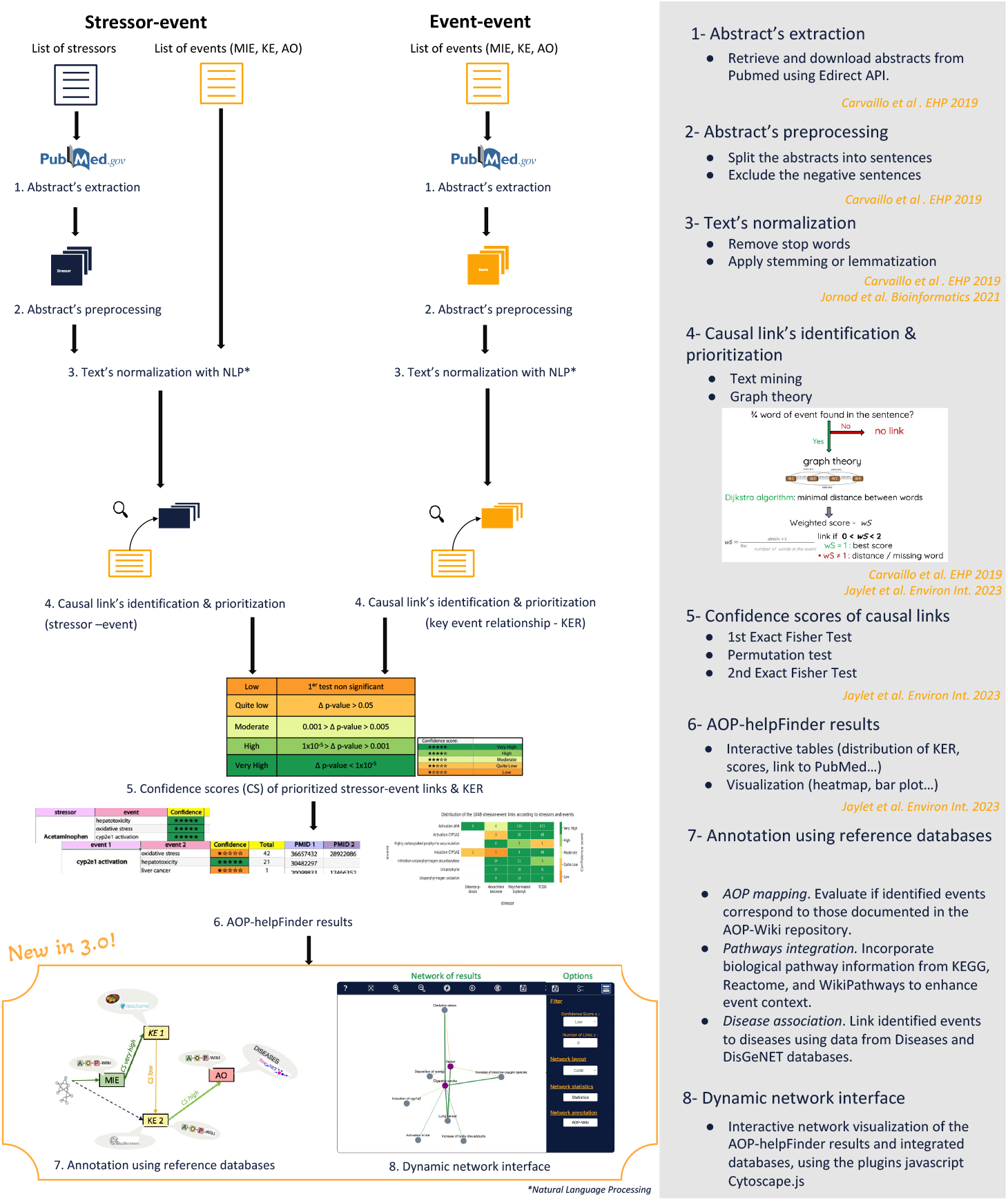
Artificial intelligence for AOP development. Workflow procedure of the AOP-helpFinder 3.0 tool.

### 2.2 Knowledge annotation using multi-source databases

Among them, the AOP repository named AOP-Wiki, which directly links the identified events to existing AOPs, was integrated (https://aopwiki.org/). Other databases provide complementary information, such as the Human Protein Atlas (HPA), which details gene expression in different tissues, various signaling pathways databases e.g., KEGG, Reactome, WikiPathways, Uniprot for human gene-pathway/process associations, and for human gene-disease associations the DISEASES (JensenLab, knowledge section) and DisGeNET (curated section) databases were included. The annotation adopts a systems biology approach, providing information across multiple levels of biological organization. The tool automatically retrieves the latest versions of the databases, except for DisGeNET, which is based on a fixed version (see supplementary materials section I and Table S1).

Since AOP-helpFinder searches for links based on user-provided inputs and given that the information in the databases may not be uniformly formatted, data standardization is performed to ensure efficient and accurate automatic annotation. This standardization is achieved through NLP methods (Python; NLTK v. 3.8.1), including stop word removal, stemming, and the consideration of synonyms (e.g., different synonyms for the same biological event or gene) (see supplementary materials section II and Figure S1).

### 2.3 Interactive visualization in the form of biological networks

AOP-helpFinder 3.0 offers visualization of all findings generated by the method as biological networks. The networks are integrated into a web page specific to each user and uses the Cytoscape.js package (v. 3.19.1) and JavaScript, allowing the opening and processing of data from a .json file containing network information (nodes, edges, descriptions, annotations).

The default settings display all associations extracted from the literature by AOP-helpFinder, with nodes representing biological events or stressors and edges representing the relationships. Edges are weighted based on the number of links and colored according to the confidence score assigned to each association (Figure S2) (see details of the confidence in Jaylet *et al*. 2023). Users can click on an edge to display a table containing all PubMed articles related to the association (Figure S3). The network also allows filtering results based on the number of PubMed links extracted by AOP-helpFinder and/or the confidence score assigned to these links (from the ‘main menu’ tab – Figure S4). Additionally, it enables displaying annotations for individual events (by clicking on a node) or for the entire network (from the ‘main menu’ tab) provided by the different databases (Figures S4 to S7).

The generated network can be saved as an image (.png or high-quality vector image .svg, ideal for publications), and users can export all information as a table (.tsv) readable in Cytoscape, allowing for more advanced analysis of the results. The network features a ‘Help’ tab (?), containing a description of all functions (Figure S2).

### 2.4 Redesign of the web server

Moreover, the AOP-helpFinder web server has been redesigned with a more modern and user-friendly interface, adding new functionalities. The homepage now offers two distinct options: (1) searching via AOP-helpFinder and (2) visualization of previous results as networks (Figure S8). The AOP-helpFinder search now includes new options, allowing users to choose between the original TM method (which requires finding at least ¾ of the event’s words to consider a link in the literature) (Jaylet et al. 2023) and a more precise but restrictive method that requires finding all words, thereby reducing the risk of false positives. The tool also allows for combined stressor-event and event-event searches, enabling the generation of a computational pre-AOP (or AOPN) and linking stressors to this pre-AOP. This TM step is then automatically complemented by annotating the biological events with database information. This search requires the user’s mail address to access the upload page and to receive notifications when results are available for download or network visualization. As with previous versions, all results and mail addresses are automatically deleted after one month, ensuring privacy and aligning with digital sobriety to reduce environmental impact by limiting computing use.

## 3 Case study: developing a computational AOPN induced by radon exposure

The set of exposures an individual encounters throughout his life is conceptualized as the exposome (Wild 2005). Many different components represent the exposome, such as chemicals, including pollutants, pesticides, additives, and drugs, as well as physical factors such as radiation emitted by radon, which can pose significant challenges to human health and ecosystems. Chronic residential exposure to radon is currently considered the second leading cause of lung cancer worldwide, after tobacco, and the primary cause among non-smokers (WHO 2009). Epidemiological studies have also suggested radon’s involvement in extra-pulmonary diseases, such as neurodegenerative diseases (Lehrer, Rheinstein and Rosenzweig 2017), stomach cancers (Barbosa-Lorenzo, Barros-Dios and Ruano-Ravina 2017), and an increased risk of leukemia (Rericha et al. 2006, Lu et al. 2020). Although the majority of inhaled radon deposits in the lungs and irradiates lung tissue, studies have shown that a small fraction can enter systemic circulation and distribute to various tissues, including bone marrow, affecting hematopoietic stem cells and potentially explaining observations regarding leukemia (Richardson, Eatough and Henshaw 1991, Sakoda et al. 2010), as well as the brain, affecting radiosensitive glial cells and contributing to neurodegenerative disorders (Zhang et al. 2022). Additionally, radon, found naturally in the environment, can also deposit in water, potentially posing a risk for stomach cancers (Barbosa-Lorenzo, Barros-Dios and Ruano-Ravina 2017). However, while residential radon exposure seems to induce extra-pulmonary effects, none of these effects have been causally demonstrated, and the mechanisms associated with these pathologies remain undefined.

To gain a comprehensive understanding of the mechanisms and risks associated with radon exposure, we took advantage of the AOP framework. We conducted a ‘stressor-event’ search using AOP-helpFinder 3.0 to automatically identify existing relationships between radon and a list of 41 biological events including genes identified by experts as potentially linked to radon (Table S2). This search was followed by an ‘event-event’ search to connect the biological events, providing a confidence score for these relationships (KERs). This TM step proposed an AOPN containing 28 of the 41 events linked to radon exposure. These events were then automatically annotated by AOP-helpFinder 3.0 using toxicological databases. Out of the 28 events, 15 were annotated, yielding more than 800 biological insights (AOPs, diseases, pathways) (examples are provided on Figure S9). The annotations produced significant results. As example, the AOP-Wiki annotation linked our events to various AOPs focusing on different cancers, including lung cancer, breast cancer, urogenital cancer, liver cancer, and leukemia, as well as other disorders such as neurological and vascular diseases, elucidating potential mechanisms. Additionally, gene annotations with KEGG, DisGENET, and DISEASES databases corroborated the same diseases and disorders, further validating our findings. Gene annotations also highlighted key biological pathways associated with radon exposure, such as cell cycle disruption, apoptosis, and transcriptional disturbances, particularly related to *TP53*, as well as other pathways also important in cancer and neurological disorders.

The network presented in Figure 2 displays the organized results as a fully computational AOPN, highlighting the most pertinent annotations for this case study. In summary, this computational AOPN, integrating literature information and annotations from databases, presents consistent results and demonstrates that radon primarily exerts its deleterious effects through DNA damage, promoting mutations (e.g., *TP53, KRAS, EGFR*) and alterations in gene expression (e.g., *CDKN2A*) involved in key pathways such as apoptosis and the cell cycle, leading to disease development. Notably, these toxic mechanisms appear to be similar across different diseases. Although these predictive results should be interpreted with caution, this case study illustrates the potential of AOP-helpFinder 3.0 to integrate relevant information from multiple sources, providing a starting point for future research and supporting modern, predictive toxicology.

**Figure 2.**
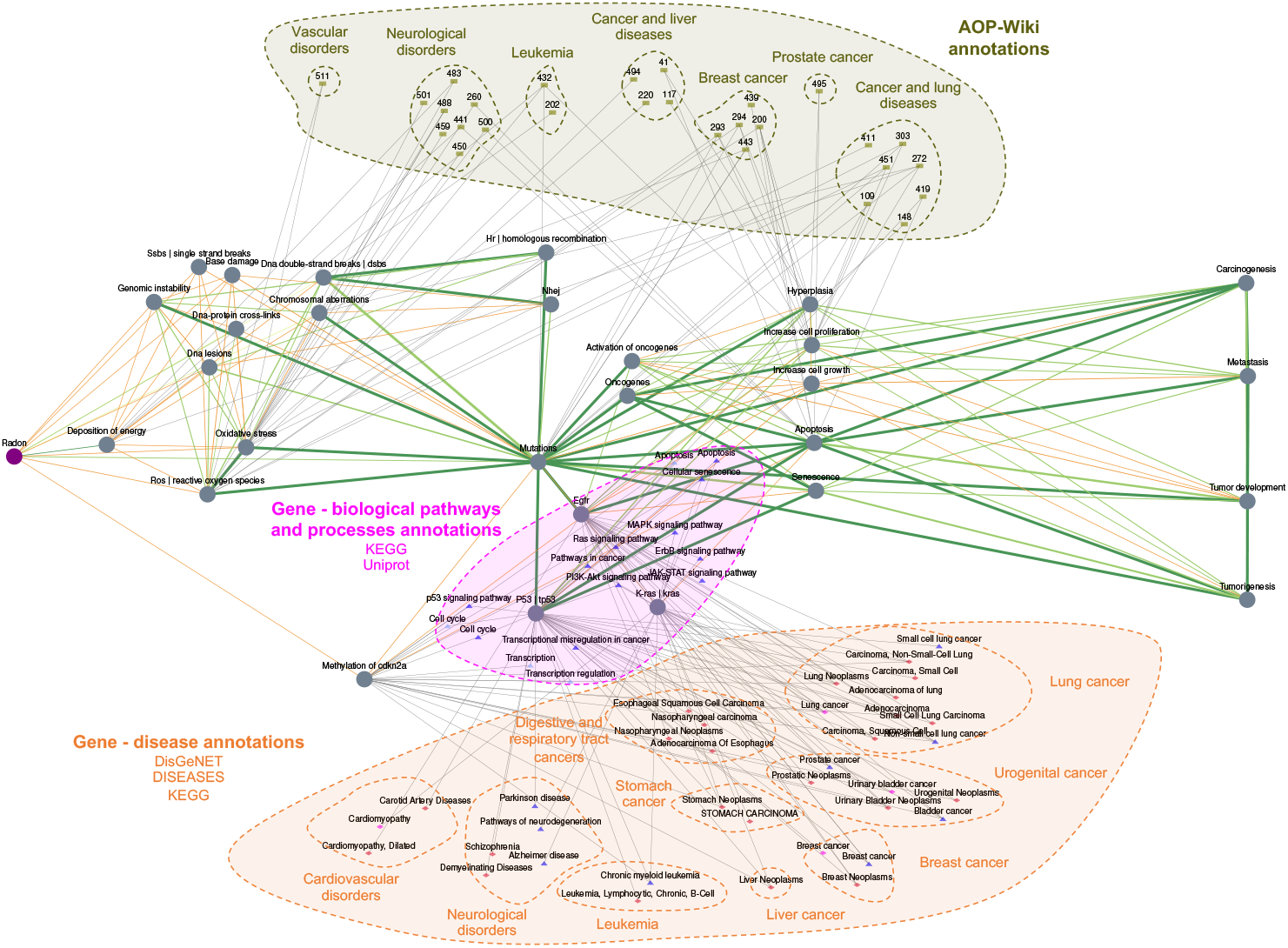
Computational AOP network initiated by Radon exposure. The round nodes represent the stressor (purple) and biological events (gray) extracted from PubMed literature using text mining, connected by edges weighted according to the number of articles addressing the link, and colored based on the confidence score (orange: Low; yellow: Moderate; light green: High; dark green: Very High). Annotations from databases are surrounded by dashed lines: AOPs in khaki, gene-disease annotations in orange, and gene-pathway annotations in magenta.

## 4 Conclusion

AOP-helpFinder 3.0 is a powerful, interactive, and user-friendly tool designed to efficiently identify, extract, and prioritize biological knowledge. By conducting an in-depth screening of scientific literature and annotating relevant information with complementary toxicology data from multiple databases, it enables rapid insights. Results can be viewed directly on the web server as comprehensive biological networks, facilitating the development of computational AOPs (Adverse Outcome Pathways) and AOPNs (AOP Networks). This tool supports next-generation risk assessment by incorporating mechanistic insights and helps pinpoint knowledge gaps, guiding the direction of future research efforts.

## Acknowledgements

The authors would like to acknowledge INSERM and the Université Paris Cité for supporting the work.

